# Tick-borne zoonotic flaviviruses and *Borrelia* infections in wildlife hosts: what have field studies contributed?

**DOI:** 10.1101/2023.08.25.554822

**Authors:** Armelle Poisson, Thierry Boulinier, Laure Bournez, Gaëlle Gonzalez, Camille V. Migné, Sara Moutailler, Bruno Faivre, Raphaëlle Métras

## Abstract

Tick-borne flaviviruses and *Borrelia* spp. are globally spread pathogens of zoonotic potential that are maintained by a transmission cycle at the interface between ticks and vertebrate hosts, mainly wild animals. Aside data on pathogen burden in ticks, information on the status of various hosts relative to infection could be critical. We reviewed how those infections have been studied in wildlife species in the field to discuss how collected data provided relevant epidemiological information and identify needs for further studies. The literature was screened for populational studies on direct or antibody detection for tick-borne *Borrelia* spp. and flaviviruses in animals, in the wild. Overall, *Borrelia* spp. were more studied (73% of case studies, representing 297 host species) than flaviviruses (27% of case studies, representing 114 host species). Studies on both *Borrelia* spp. and flaviviruses focused mainly on the same species, namely bank vole and yellow-necked mouse. Most studies were order-specific and cross-sectional, reporting prevalence at various locations, but with little insight into the underlying epidemiological dynamics. Species with potential to act as reservoir hosts were overlooked, notably passerine birds. We highlight the necessity of collecting both demographics and infection data in wildlife studies, and to consider communities of species, to better estimate zoonotic risk potential in the One Health context.

## I Introduction

Vector-borne and zoonotic infections are emerging threats to public health[1]. Tick-borne pathogens (TBP) are maintained by multi-hosts complex epidemiological cycles[2]. Transmission to humans occur on specific occasions, and understanding and quantifying it is necessary to control diseases. Persistence of tick-borne and zoonotic pathogens is allowed at the interface between tick vectors and a large variety of vertebrate hosts permitting ticks to feed and complete their life cycles. In those systems, early tick stages (larvae and nymph) usually feed on small mammals and birds, while mature stages mainly feed on larger vertebrates (such as deer, boar, cattle)[3].

Beyond their feeding role for ticks, vertebrate species can act as pathogen reservoir or sentinel hosts. Infection and exposure data from both give precious information on dynamics of the studied TBP[4]. Whilst reservoir hosts allow the pathogen to multiply and be transmitted further, sentinel species are characterized by their ability to reflect underlying epidemiological phenomena and by being of easy access[5]. For instance, wild red and Arctic foxes have been suggested as sentinels for human and animal toxoplasma risk in Canada, based on serology and direct detection on carcasses[6]. Also, animal ecology can drive variations in risk of transmission to human[7]. Therefore, because understanding pathogens’ transmission and distribution in wildlife is prerequisite to addressing risk of transmission, quantitatively tracking infection and exposure to pathogens in those hosts is necessary.

Tick-borne *Borreliaceae* and flaviviruses are TBP of zoonotic potential which are the most prevalent in the temperate regions of the world[8], and are thus of interest for human and veterinary public health. Tick-borne encephalitis virus (TBEV), first described in Austria in 1931, is the most common tick-borne flavivirus (TBFV) in Europe with several endemic foci in Europe and Asia and growing number of human cases[9]. Its ecology and epidemiology have early been studied in Europe, especially in Czech Republic[10]. Other TBFV have later been isolated from wild animals and their ticks, such as Meaban virus (MEAV) in Brittany, France[11], or from human cases, such as Powassan virus (POWV) in Powassan town, Canada[12]. Prominent vectors of TBFV are hard ticks *Ixodes ricinus* in Western and Central Europe and *I. persulcatus* in Eastern Europe, as well as local tick species in other parts of the world. Similarly transmitted bacterial genospecies of the *Borrelia burgdorferi* sensu lato (Bbsl) complex are the causative agents for Lyme disease, one of the most common TBD in the Northern Hemisphere. Other tick-borne *Borrelia* spp. cause relapsing fever in humans[13].

Recent reviews on tick-borne *Borrelia* spp. and TBFV have focused on getting information on the biology of infection in vertebrate hosts[14,15], reporting prevalence and clinical cases in human[16,17] and discussing diagnostic methodologies[18,19], the role of non-vector transmissions for TBFV[20,21], or on the importance of modelling TBP regarding climate change[22,23]. Some reviews have focused on wildlife hosts but only at a national or continental scale[24,25], or on reservoirs associated with flaviviruses in general, with little focus on TBFV[26]. Finally, a lot of research effort has focused on studying the infectious agents in the ticks[27,28].

Despite their major importance in zoonotic transmission, no study has reviewed the body of evidence in wildlife hosts from field data at a global scale. Yet, data on the level of infection and exposure in free-ranging populations are critical to model the dynamics of infectious diseases[29]. Strong inference about the transmission processes underlying those dynamics often requires a combination of approaches, notably experiments to ascertain the role of reservoir[30], but data on infection and exposure of hosts are prerequisite in most cases. When it comes to wildlife species, a strong heterogeneity in the type of field data is nevertheless expected, from local cross-sectional sampling of particular species, to broad spatial surveys over series of years. Estimating epidemiological parameters also requires accounting for potential biases in detection probabilities and host population parameters[31], which demands specific designs.

The aim of this review was thus to investigate how tick-borne *Borrelia* spp. and flaviviruses infection burdens have been studied in free-ranging wildlife (that is, excluding studies of TBP in ticks) to discuss how collected data provided relevant epidemiological information, and identify needs for further studies. We predicted that strong heterogeneity would appear in the types of hosts investigated, with a high number of studies conducted on common species sharing landscape use with humans in the Northern Hemisphere. We expected few studies including whole communities of potential hosts and few studies estimating TBP burden while accounting for temporal and spatial variations and uncertainties. We anticipated that monitoring of wildlife diseases would have been difficult and that knowledge of what is precisely measured would have been uncertain.

### II Studied hosts and their potential role in epidemiological cycles

We reconstructed 947 “case studies” (unique combinations of species-pathogen-area-study period) studying overall 347 host species, from the 314 articles matching the selection criteria (Figure 1). Detailed methodology on the literature search and metrics on the results are available in SuppInfo1. Figure 2 links the order of the tested species to the investigated pathogen.

**Figure 1:**
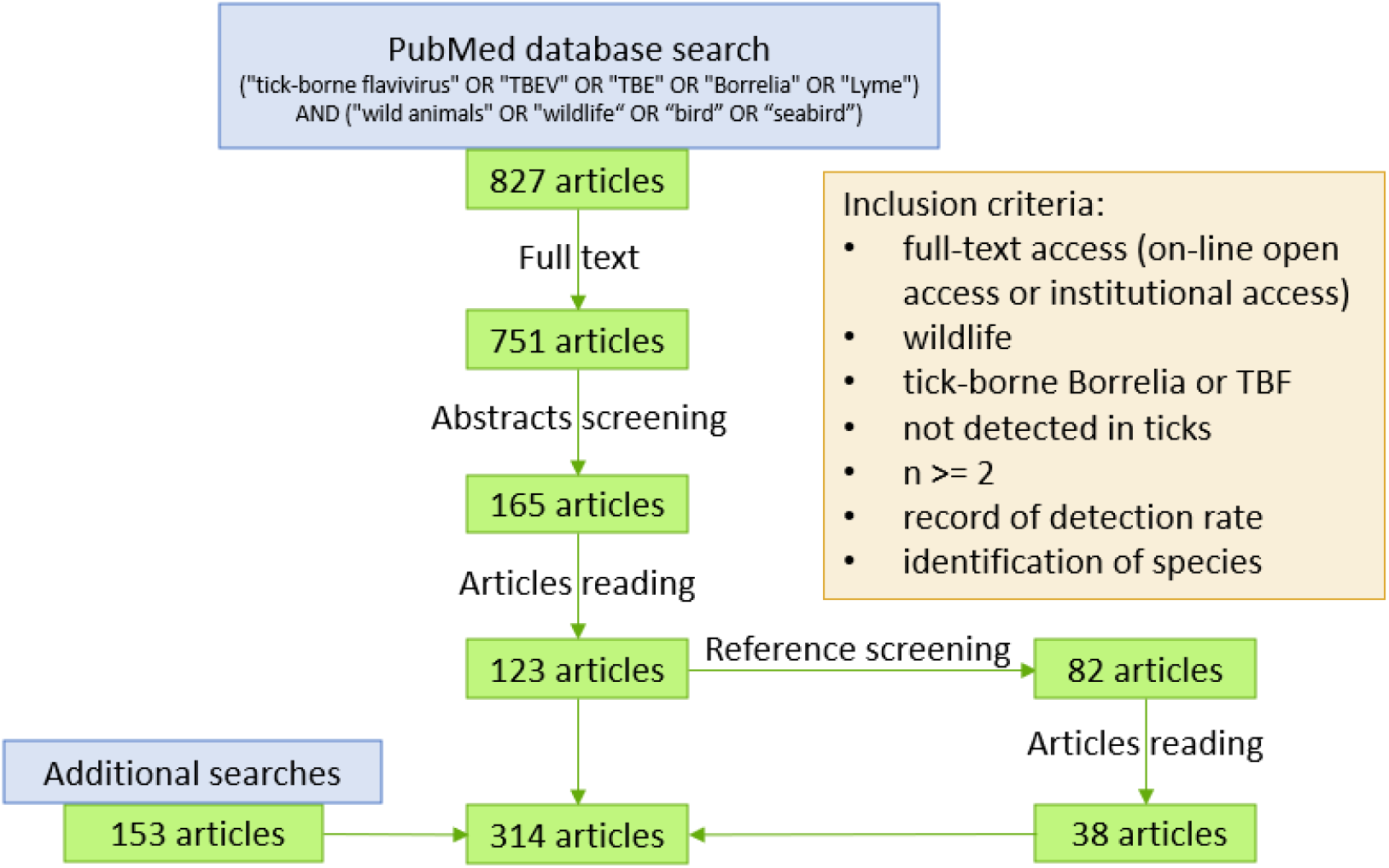
Diagram of inclusion and exclusion in database screening. See SupInfo1 for details on literature search and additional searches.

**Figure 2:**
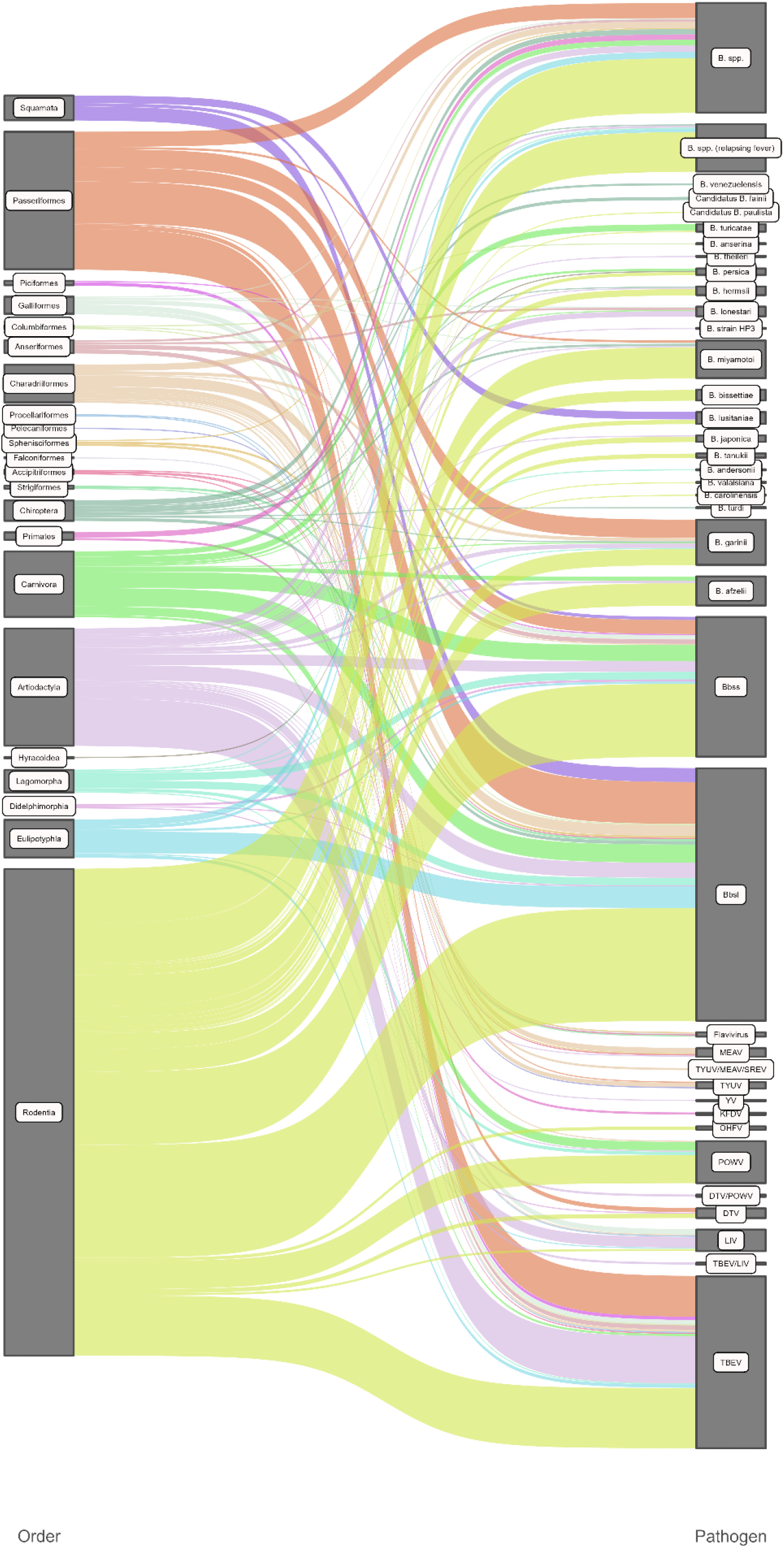
Sankey diagram pairing study cases by pathogen and order. Width of the ribbon is proportional to the number of cases included in this order-pathogen couple. Data are available in Supplementary Dataset 1.

### a Historical papers shaped research on rodents as reservoirs

Rodentia was the most investigated order, represented by more than half (363/689) of the *Borrelia* study cases and more than a third (91/258) of the TBFV study cases. White-footed mouse (*Peromyscus leucopus*) was investigated in the first published studies from Northern America, and has been regarded since the eighties as the most common *Borrelia* spp. reservoir host in wildlife in Northern America[32]. Regarding TBFV, a series of studies were conducted initially in Northern America too. The first published studies looked at Powassan virus (POWV) in *Peromyscus* genus, only present in Northern America, and secondly in *Microtus* and *Mus* genera, present in the continent as well. In addition, in Canada, a series of studies from the same authors, presented rodents as primary reservoirs for POWV. For this, species such as *Tamias, Urocitellus* and *Callospermophilus* chipmunks and *Tamiasciurus* and *Sciurus* squirrels were investigated, with average seroprevalence of 15%[33–41]. Groundhogs (*Marmota monax*) sampled in the same studies reported the highest seroprevalence (around 43%)[37–39]. In parallel, several studies on the eco-epidemiology of TBFV were conducted in the former Union of Soviet Socialist Republics but were not published in English, although some articles are available with translation systems such as those of NAMRU-3 (see limitations in SuppInfo1). Rodents were later sampled on other continents and TBFV detections were several times successful in Europe in *Apodemus, Microtus* and *Myodes* genera[42], and in *Micromys, Mus* and *Sciurus* for *Borrelia*[42,43]. In Europe, the bank vole (*Myodes glareolus*) and yellow-necked mouse (*Apodemus flavicollis*) were recognized as reservoirs for TBEV[44]. Bbsl was detected in *Tamias, Tamiasciurus* and *Neotoma* in North America[45]. Furthermore, *Gerbillus, Mastomys, Praomys* and *Apodemus, Cricetulus, Eothenomys, Meriones, Mus, Niviventer* and *Rattus* bore relapsing fever associated *Borrelia* respectively in Africa and in Asia. Notably, Eastern grey squirrels (*Sciurus carolinensis*) similarly carried *Borrelia* DNA, both in their native countries and where they are invasive in Europe.

### b Deer, broadly studied as sentinels

The second most investigated order by the number of tested individuals was Artiodactyla. Artiodactyls were mainly represented by the *Cervidae* family, in terms of numbers of tested animals (82%, 26,090/31,862) and investigated species (56%, 9/16) (Figure 3B and 3D). Deer are considered incompetent reservoirs for *Borrelia* spp. by some authors because of borreliacidal properties of their sera[46]. Yet, studies showed that the complement borreliacidal activity depends on strain and also exists on other *Borrelia* strains for other vertebrates, including rodents[47]. Despite their poor competence as reservoirs, deer are frequently met across the world and can be infected by feeding adult ticks they commonly host[48]. In *Cervidae*, species locally present in America, Europe and Asia showed similar level of TBP contact, with average seroprevalence of 18%. Restraint home ranges, of no more than 10km2 among the different deer species[49,50], allowed investigating infection processes at small spatial scales. Moreover, surveillance activities and pathogen detection can be facilitated through deer hunting, allowing widespread sampling programmes on a regular basis for reasonable costs and working efforts[51]. As such, deer could help identifying endemic or emerging risk areas and give information on underlying processes, and were repeatedly considered as sentinels. Other game artiodactyls identified in this review, such as wild boar (*Sus scrofa*), offer the same kind of possibilities for sentinel surveillance.

**Figure 3A-D:**
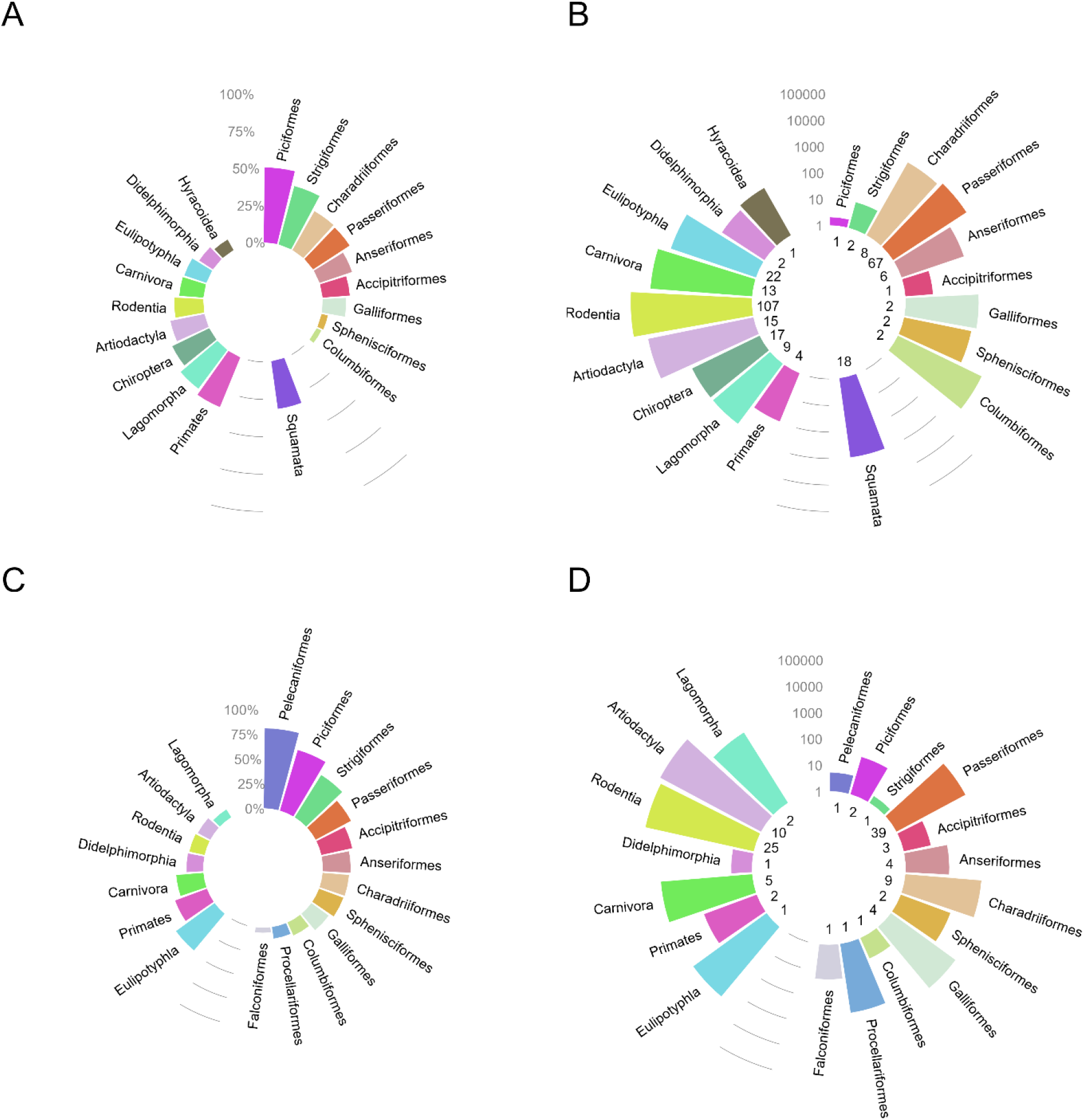
Investigation efforts for TBP by Orders. Both serological and “active infection” data are presented on the same plot. Top row shows for *Borrelia* spp. **(A)** the average percentage of animals positive to *Borrelia* by order, **(B)** the number of tested animals in each order. Bottom row shows for TBFV **(C)** the percentage of animals positive to TBFV by order, **(D)** the number of tested animals in each order. Most publications regarding TBFV used an indirect methodology (antibody detection), direct detection was used in only 11% of TBFV cases. **(B)** and **(D)** show the number of tested individuals on a log10 scale, and the number of different host species tested in each order is shown at the bottom of each bar. Orders are grouped by class, Mammalia are shown to the left, Aves to the right, and for *Borrelia* spp., Lepidosauria at the bottom. Data are available in Supplementary Dataset 1.

### c Birds, reservoirs, have been overlooked

Birds were also reported as reservoirs in the literature[52]. Contrary to rodents, birds may have been overlooked. Indeed, although birds represented about 19% (129/689) of the *Borrelia* study cases (Figure 2) and 33% (85/258) of TBFV cases, they originated from only 12% (28/228) and 18% (16/89) of the publications. In those, pathogen presence was often screened in birds caught in a banding configuration. For instance, in one study, Newman *et al*. caught birds on the University of California’s campus in 2003-2004[53]. Their wide-range study allowed detecting Bbsl in 23 species out of 53 caught species. Mistnetting is a common field method efficient for capturing birds, hence for detecting potential reservoirs. However, this approach is mostly favourable for Passeriformes, and passerines were the avian order the most often studied as potential reservoirs for TBP. In addition, for migratory species or those exploiting large areas during their life cycle, the infection status might not always reflect the local infection processes. Nevertheless, combined with movement information, they can unravel critical information on pathogen dissemination over long distances and between various ecosystems[54]. We hypothesized that limited availability of data on birds might be due to logistical difficulties related to blood sampling in the wild (training and legal requirements, ethical concerns for endangered species, practical implementation). However, antibodies against Gadgets Gully and Saumarez Reef viruses were extensively studied in a collection of Australian seabirds’ sera[55]. This constitutes a unique occurrence of exposure information on rare TBFV, but sadly, no clarification was made on which species retrieved seropositivity.

### d Least studied species with growing interest

Other vertebrate species of interest have been less studied. Recently, however, new species of tick-borne *Borrelia* spp. have been sought and encountered in vertebrates from other orders. For example, squamates were investigated for *Borrelia* in America, Africa and Europe, and 18 species associated with Bbsl. Anti-Bbsl antibodies were detected in raccoon (*Procyon lotor*) in the USA and raccoon dog (*Nyctereutes procyonoides*) in Asia, as well as where they are invasive in Europe or Asia. In India, primates were found seropositive to KFDV, a TBFV close to POWV and DTV[56]. In the UK, an isolated clinical case of lethal borreliosis was reported in a *Pipistrellus* spp. bat[57]. Neotropical bats have been identified to carry particular strains of relapsing fever *Borrelia* and Bbsl in Central and Southern America. Studies in Central America have been conducted as follow-back of a paper published in 1968 by Marinkelle & Grose[58]. Some areas are currently investigated more in depth or monitored. For instance, Tribeč mountain region in Czech Republic is continuously investigated for TBEV since the seventies.

## III Diagnostic tests: current *versus* past infections

Direct and indirect detection methods were at different use depending on the pathogen. Whilst most *Borrelia* studies (71%) looked for pathogen’s genome with molecular techniques to identify ongoing infections, 89% of flaviviruses studies used serological methods. Long-lasting in the organisms, *Borrelia* spp. were often sought by PCR (60% of case studies overall). Skin and blood were used for spirochetes PCR detection, allowing positive samples to be sequenced and species identified, although the use of blood compared with other organs to directly detect *Borrelia* is discussed[59]. Determination of *Borrelia* subspecies was useful in understanding the role of hosts in disease cycles[60]. Different *Borrelia* have been preferentially detected in different hosts, such as *B. lusitaniae* which was the only Bbsl genospecies found in lizards in Europe.

In contrast, the short viremia noted after flaviviruses infection as demonstrated by experimental infections[61], makes direct viral detection challenging especially in wildlife. PCR was therefore rarely used for flaviviruses (in only 11% of TBFV studies), but allowed sequencing of TBEV subtypes that clustered with TBEV-Eur (European) in Croatia, Slovakia and the Netherlands[62–64], TBEV-Eur and TBEV-Sib (Siberian) in Finland[65], TBEV-Eur, TBEV-Sib and TBEV-FE (Far Eastern) in Russia[66] and TBEV-Him (Himalayan) in China[67], and evidenced co-infection in Russia[66]. Identifying flaviviruses will help understanding further the ecology and dynamics of pathogens and will assist disease surveillance with phylogenetics providing information on circulation and evolution of TBP[68]. For example, TBEV could be mistaken for Looping Ill virus (LIV) causing severe conditions mainly in sheep[51].

Flaviviruses infections induce immune response with neutralizing antibodies that could be detected lifelong in blood in human[69] and up to 168 days after experimental infection in bank voles[70]. ELISA were the most used serological techniques (89% of flaviviruses cases, and 38% of overall cases). Serological information in combination with metadata can provide precious information on pathogen transmission[71]. Yet, cross-reactivity in wide-range immunoassays may hamper data interpretation[72]. For instance, anti-MEAV antibodies were surprisingly detected in *Cervidae*[73], whilst MEAV belongs to the seabird group in TBFV phylogeny[74]. Antibodies against LIV have been unusually identified in seabirds *Fratercula arctica* and *Hydrobates leucorhous*[75]. In addition, whilst ELISA cut-off is generally reproduced from manufacturer, some authors proposed its determination via mathematical fit on observed data to optimize sensitivity and specificity[76]. Others even chose to present raw ELISA results as a signal of contact with TBFV, without indicating cut-off nor prevalence[77].

Publications about TBFV outside of TBEV and POWV were marginal and specific detection for these TBFV antibodies was not always commercially available. Arnal *et al*. found MEAV-seropositivity in *Larus michaellis michaellis* egg yolks in France using flavivirus-ELISA cross-reactivity properties, with the ID-Vet ID Screen® kit designed for the mosquito-borne flavivirus West Nile virus (WNV)[78]. Similarly, through both COMPAC® WNV-ELISA and exclusion via serum neutralization tests (SNT), Jurado-Tarifa *et al*. investigated flaviviruses in Spanish birds of prey and detected evidence of exposure to flaviviruses different from the mosquito-borne WNV and Usutu (USUV) viruses and from MEAV in *Circus pygargus, Falco tinnunculus, Hieraaetus pennatus* and *Tyto alba*[79]. This relies on lack of specificity of commercially available anti-WNV antibodies detection kits. Some commercial TBEV ELISA have also been compared for their reliability according to the TBEV-strain used and for the at-risk cross-reactivity[80]. In the UK, Holding *et al*. recorded similar LIV and TBEV seroprevalences in cervids although TBEV was though not to be present on the British Isles[51]. Holding et *al*. used the FSME/TBE All-Species Progen® ELISA and a hemagglutination assay specific to LIV. For confirmation, Holding *et al*. sequenced TBEV in a tick itself. Ytrehus *et al*. also showed presence of both LIV and TBEV antibodies in Southern Norway in *Alces alces* and *Capreolus capreolus* using ELISA kits for TBEV and hemagglutination for LIV as well as SNT for verification[81]. However, their results appeared differentiable for the two pathogens. On the opposite, Bournez *et al*. chose to deliver their final TBEV micro-neutralization test serology results under the TBEV/LIV label, after a first antibodies screening in *Capreolus capreolus* and *Sus scrofa* with the ID-Vet ID Screen® WNV-ELISA[82]. TBEV and LIV as well as the Turkish Sheep Encephalomyelitis virus (TSEV) and the Spanish Sheep Encephalomyelitis virus (SSEV) belong to the TBEV serocomplex[56], hence can particularly easily be mistaken in serology.

## IV Infection and exposure in hosts: summary measures

A total of 297 wild animal species (mammals, birds and lizards) were investigated for *Borrelia* (Figure 3A and 3B). When considering all *Borrelia* species, all studies confounded, overall detection and seroprevalence tended to uniformize between 15% and 30% as the number of tested individuals increased (Figure 4A). In studies with over 50 tested individuals, the proportion of active infections ranged from 0.1% for *B. lusitaniae* in Eastern grey squirrel (*Sciurus carolinensis*) to 76% for Bbsl in white-footed mouse[83], and from 0.2% for relapsing fever associated species in house mouse (*Mus musculus*)[84] to 50% for *B. miyamotoi* in wild turkey (*Meleagris gallopavo*)[85]. In mammals, including rodents, artiodactyls and carnivores, the level of direct detection and seroprevalence of *Borrelia* spp. was between 20 and 25% for orders with over 100 specimens tested (Figure 3A and 3B).

**Figure 4A-B:**
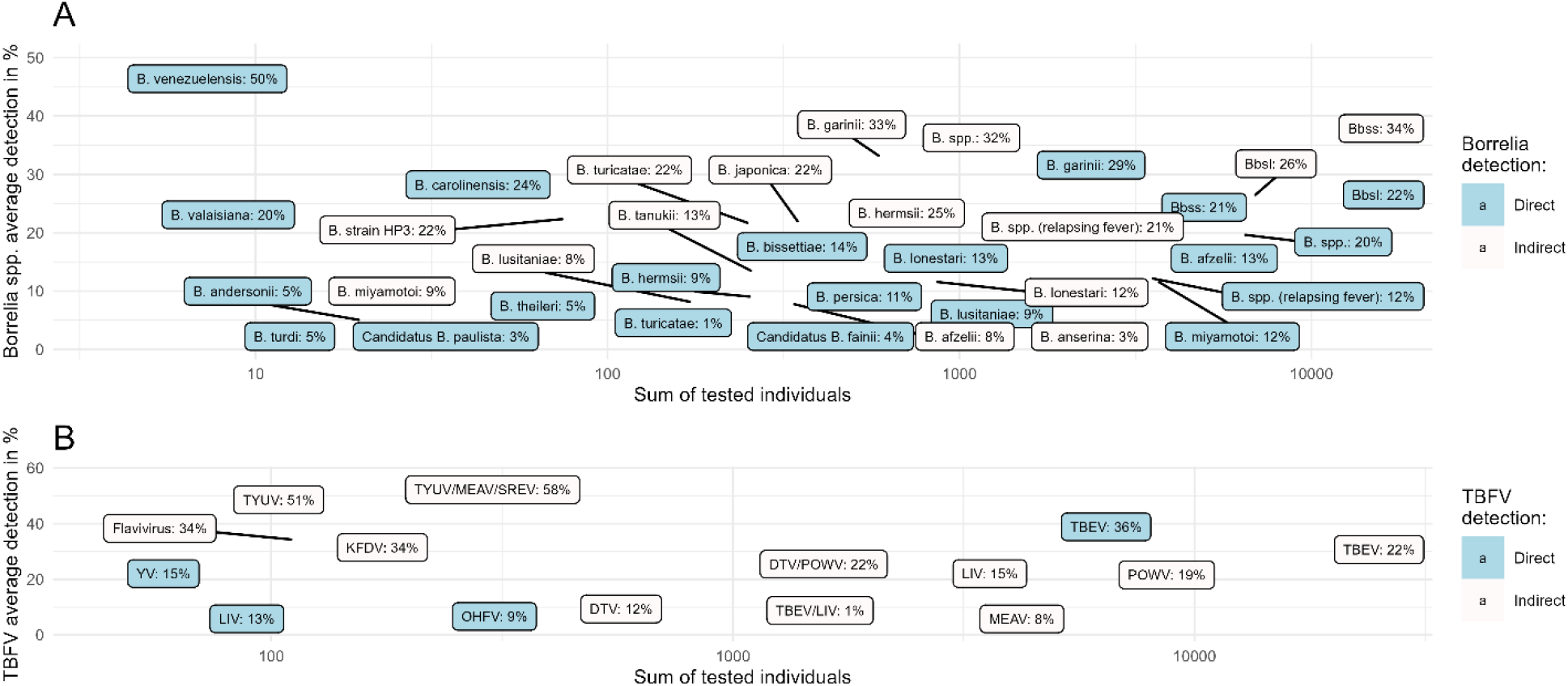
Average prevalence of pathogen detection (dark blue labels, direct tests) and antibodies (light grey labels, indirect tests) in the tested individuals, rounded to 0 digit. The x-axis represents the log10 of the sum of individuals that have been tested for **(A) *Borrelia* spp**. and **(B) Flaviviruses** in the selected publications, the y-axis represents the % of positive. DTV (Deer Tick Virus) in oldest publications might have been detected under the name of POWV, because DTV formerly corresponded to POWV lineage II. Data are available in Supplementary Dataset 1.

In rodent studies testing more than 50 individuals, the average proportion of *Borrelia* direct detection was 11.2% (median 5.3%, range 0.2%-76%) and average seroprevalence was twice higher (mean 21.2%, median 15%, range 0.7%-89%) (Table 1, Figure S2). Variations were reported along study sites and periods regardless of taxonomy. Beyond that such TBP might be widespread, this could be explained by the ecology broadly shared by rodents, living close to the ground where foraging infected ticks can be encountered[86], and therefore leading to similar exposure. Variations in prevalence across studies were observed notably for a same species, such as for Bbsl direct detection in yellow-necked mouse tissues varying from 5.8% in Hungary in 2014-2015[87] to 23% in Germany in 2012-2014[88]. In artiodactyls studies with more than 50 sampled animals, the average prevalence of *Borrelia* DNA detection was 14.4% and average seroprevalence was 23% (Table 1, Figure S2). They were of 12.6% and 24.2% in birds (Table 1, Figure S2). Among birds, Columbiformes, Passeriformes, Charadriiforms and Galliformes were the most studied in terms of number of tested individuals, with Charadriiforms and Passeriformes reaching 29% of seroprevalence (Figure 3B). Among the least studied species, Chiropters presented both seropositivity (36%) and pathogen prevalence (25%) of rather high level (Figure 3A).

**Table 1:** Prevalence of direct detection and seroprevalence by Order, extracted from studies with more than 50 individuals. n = number of study cases, median values, mean, range and inter-quartile range (IQ). See plots of distributions in Figure S2.

| Studies with >=50 tested individuals in: |  | Rodents (n=230) | Artiodactyls (n=83) | Birds (n=55) |
| --- | --- | --- | --- | --- |
| ***Borrelia*** | **Direct detection** | n = 135;<br>median 5.3%;<br>mean 11.2%;<br>range 0.2-76%;<br>IQR 2-15%; | n = 15;<br>median 10.6%;<br>mean 14.4%;<br>range 0.8-49.7%;<br>IQR 6.4-17.4%; | n = 22;<br>median 8.8%;<br>mean 12.6%;<br>range 1-51%;<br>IQR 4.4-14%; |
|  | **Seroprevalence** | n = 43;<br>median 15%;<br>mean 21.2%;<br>range 0.7-89%;<br>IQR 9.2-31.5%; | n = 25;<br>median 19%;<br>mean 23%;<br>range 0.8-68.8%;<br>IQR 7.2-30%; | n = 14;<br>median 17%;<br>mean 24.2%;<br>range 3-78.6%;<br>IQR 9.5-26.6%; |
| **TBFV** | **Seroprevalence** | n = 44;<br>median 5.0%;<br>mean 8.9%;<br>range 0.4-44%;<br>IQR 3.5-10.5%; | n = 42;<br>median 5.4%;<br>mean 10%;<br>range 0.07-62.7%;<br>IQR 2.2-10%; | n = 18;<br>median 19.7%;<br>mean 21.9%;<br>range 0.5-68.1%;<br>IQR 4.1-33.8%; |

TBFV or antibodies were searched and detected in 114 mammal and bird species (Figure 3C and 3D). The overall average seroprevalence ranged from 8% for MEAV to 51% for TYUV (Figure 4B), but with heterogeneous sample sizes. Among studies with more than 50 tested individuals, seroprevalences ranged from 0.07% for MEAV in red deer (*Cervus elaphus*)[73] to 68% for TYUV/MEAV/SREV-like in European herring gull (*Larus argentatus*)[89]. For TBFV of the mammalian group, seroprevalence ranged from 0,2% for TBEV/LIV in roe deer (*Capreolus capreolus*)[82] to 63% for TBEV in European bison (*Bison bonasus*)[90]. In rodent studies with at least 50 samples, the average TBFV-seropositivity was 8.9% (median 4.95%, range 0.4-44%) (Table 1, Figure S2), lower than for *Borrelia*. No studies looking at rodent host exposure to Kyanasur Forest Disease virus (KFDV) and Yamaguchi virus (YV) were recorded, although they have terrestrial hosts[91]. The average seroprevalence in artiodactyls was 10% and more than twice in birds reaching 21.9% (Table 1, Figure S2). Most studied avian orders were Passeriformes, Charadriiforms, Galliformes and Procellariforms, with highest seroprevalence for Passeriformes (about 40%), followed by Charadriiforms (27%) and Galliformes (21%) when Procellariforms tended to have less seropositive individuals (Figure 3C). Whilst 80% of Pelecaniformes had antibodies to TYUV, for this order only 8 European shags (*Phalacrocorax aristotelis*) were tested on a single location (in Brittany, France) in the eighties[89], highlighting the weight small scale studies could have. Exceptionally, mammalian Eulipotyphla were well-studied by PCR techniques and showed high rates of viremia (of about 50%)[92].

## V Study designs and eco-epidemiological parameters

Implemented study designs varied in spatial and temporal dimensions, resulting in varying quality in data acquisition. Most publications conducted cross-sectional studies (57%, 180/314), and 20 did not inform a study period. Some studies conducted repeated sampling without taking into consideration the sampling period in data presentation. Spatial resolution varied from locality to country scale, with often provinces chosen as spatial scale. Inconstancy came from different focuses, either on locally targeted area or over broader-scale exposure. In each study, sample sizes were usually limited, with median sample size at 40 (IQR 14-119). Overall, the way data were collected did not allow taking geography as well as other confounding factors into account. Forty-one (41) publications marked captured individuals but marking techniques were not always specific and trustable, some authors simply relying on scars made by biopsy or on feather clipping. The aim of marking was sometimes only to avoid resampling, with immediate release in case of recapture. However, 24 publications used individual marks and repeated capture occasions. Nevertheless, capture-mark-recapture data was not systematically analysed using relevant techniques. It is however an important specific sampling approach that permits accounting for detection uncertainty. For instance, seroprevalence and capture-mark-recapture data were recently used in a *Borrelia* host system[93]. Similar approaches were also used in other wildlife and infection systems, such as brucellosis in Alpine ibex (*Capra ibex*)[94].

Moreover, metadata informing basic demographics were lacking in almost all studies whereas data on age structure can be key in understanding susceptibility to pathogen at the individual level[95], but also transmission processes at the population level. Mammals and birds or squamates were not studied together in the same areas. Designs tended to investigate a single order by setting field material only fit for this order, whilst theoretical definitions of reservoir and sentinel suggest that reservoirs are communities rather than species[30]. The underlying community of hosts was seldomly considered. Furthermore, although sometimes convenient, getting material from hunted, road-killed or rescued animals implies biases in detection that should be accounted for. Such sampling strategies often lead to small sample sizes and difficult interpretation. Moreover, as recommended by Yoccoz *et al*. on biodiversity monitoring, sampling designs and data analyses need to account for uncertainty on host detectability and spatial variability, for better estimating the state and rates of change of relevant ecological communities[96]. Thereby, acquiring infection and exposure data in wildlife should target communities of species in longitudinal studies with capture-marking-recapture or patch-occupancy modelling.

## VI Complementarity of laboratory and tick data

In our review, some infectious agents received less attention, such as TYUV, MEAV and SREV seabird flaviviruses. Whilst such agents may be less pathogenic, they can still be useful to understand transmission processes. In addition, their virulence could vary between host species and they could emerge as possible pathogens of concern. For example, TUYV was unexpectedly found pathogenic and deadly in seabirds in a study conducted by Berezina *et al*. in 1974[97]. Therefore, combining data on pathogenicity from the field and the laboratory is still needed.

In the literature, models and experimental infections have attempted to characterize reservoir competence[97]. Especially, xenodiagnoses are used to detect host reservoir capacities and tick vector competences in experimental settings[98]. Ticks from the field have been extensively examined for pathogens. From the 827 articles matching the literature search, 224 publications were directly looking into ticks for pathogens with methods ranging from PCR to metagenomics. Studies on ticks can be complementary to data from hosts insofar as they analyse tick-host interactions. Their deployment is less invasive on hosts than blood or tissue sampling. They need less expertise in the field and are subject to less strict ethical protocols. Such studies can serve as early detection methods when the pathogen is poorly known. For instance, whilst data on TBP hosts in Australia were limited, studying pathogens in ticks demonstrated circulation of *Borrelia* in squamates and monotremes[99]. In addition, identification of ticks’ life stages allows to infer ticks’ contamination capacity, although other ways of transmission such as vertical transmission in vertebrate hosts have been reported for some agents[100]. However, accounting for the whole local host community and their infection status remains as important.

## VII Conclusion

TBP are present in a wide range of wild animal species, and rodents and artiodactyls were the most investigated. Some infectious agents, rare TBFV or particular *Borrelia* strains, were little studied and information on their infecting capacities is still needed. Diagnostic detection methods varied with the targeted pathogens. Selected publications showed that study designs were heterogenous, both in data collection and analysis. Key information on host age and precise location was often unavailable; 58% (200/342) of the studies conducted fieldwork at one time point, and very few studies used quantitative methods to analyse collected data. Quantifying transmission necessitates gathering empirical data from populational studies, including hosts demographics and identifying which species or community act as reservoirs and sentinels. Whilst we acknowledge that gathering this information is difficult, this review highlights the necessity to estimate eco-epidemiological parameters in wildlife infection studies. The subject discussed in this review concerns a multidisciplinary scientific community (ecology, epidemiology, microbiology, parasitology, modelling, public health), emphasizing that getting and interpreting zoonotic infection data from wildlife in a rigorous way is important to build workable One Health programmes.

## Data availability

The data that support this study are available as Supplementary Dataset 1. Source data are provided with this paper.

## Acknowledgements

This work was funded by Agence Nationale de la Recherche (ANR) under the ANRJCJC MoZArt project, grant number ANR-22-CE35-0003. Support is also acknowledged from ANR ECOPATHS project, grant number ANR-21-CE35-0016, and from IPEV ECOPATH-1151 project.

## Author information

### Contributions

AP, TB and RM conceived the paper. AP conducted the analysis and designed the figures. AP and RM wrote the paper. TB, GG, CM, LB, BF and SM revised the paper. All authors approved the final version of the paper.

